# An Envelope Protein of a Prokaryotic Organelle Function towards Conserving Catalytic Activity of the Native Enzyme at Higher Temperatures

**DOI:** 10.1101/2020.02.18.954511

**Authors:** Gaurav Kumar, Naimat K. Bari, Jagadish P. Hazra, Sharmistha Sinha

## Abstract

A classic example of an all-protein natural nano-bioreactor, the bacterial microcompartments are a special kind of prokaryotic organelles that confine enzymes within a small volume enveloped by an outer layer of shell proteins. This arrangement provides conditional metabolic aid to the bacteria. The outer shell allows selective diffusion of small molecules and sequesters toxic metabolites. In this work we use 1,2-propanediol utilization microcompartment as a model to study the effect of molecular confinement on the stability and catalytic activity of native enzymes in microcompartment. We observe a 50% decrease in the activity of free enzyme PduCDE at 45°C, while PduMCP retains its optimum activity till 50°C followed by more than 40% reduced activity at 55°C. PduBB’, the major component of the outer shell contributes to the increased catalytic activity of PduCDE. PduBB’ also prevents the unfolding and aggregation of PduCDE under thermal stress. Using a combination of experimental and theoretical studies we probe the interactions of the shell proteins PduBB’, N-terminal truncated PduB and single mutant PduB’M38L with PduCDE. We observe that all the three variants of PduB* shell proteins interact with the enzyme in vitro, but only PduBB’ influences its activity and stability, underscoring the significance of the unique combination of PduB and PduB’ in PduMCP assembly.

## Introduction

Complex living cells have intracellular organelles within which, they carry out specific life processes. The ability to restrict macromolecules to a well-defined space and volume to carry out life processes efficiently, is a characteristic feature of living things. Many bacterial species organize tiny compartment like structures with unique functions that help them to survive under adverse conditions (1,2). Studying the structure and function of such sub-cellular organizations may not just provide us with vital information regarding pre-biological compartmentalization (3) but will also pave the way for the development of synthetic reactors for future applications (4). In this aspect, bacterial microcompartments (MCP) (5) have stood out as the most sought after candidate due to their interesting features. They are exclusively made up of proteins with an envelope of shell proteins that wraps a cascade of enzymes within. Among all the MCPs discovered so far, 1,2-propanediol utilization microcompartment (PduMCP), encoded by the *pdu* operon with 23 genes is the most complex yet most studied **(Fig. 1a)** (6-10). Over the years, it has gained world-wide attention due to its significant role in the pathogenicity of bacteria like *Salmonella enterica* during inflammatory diseases (11).

**Fig 1:**
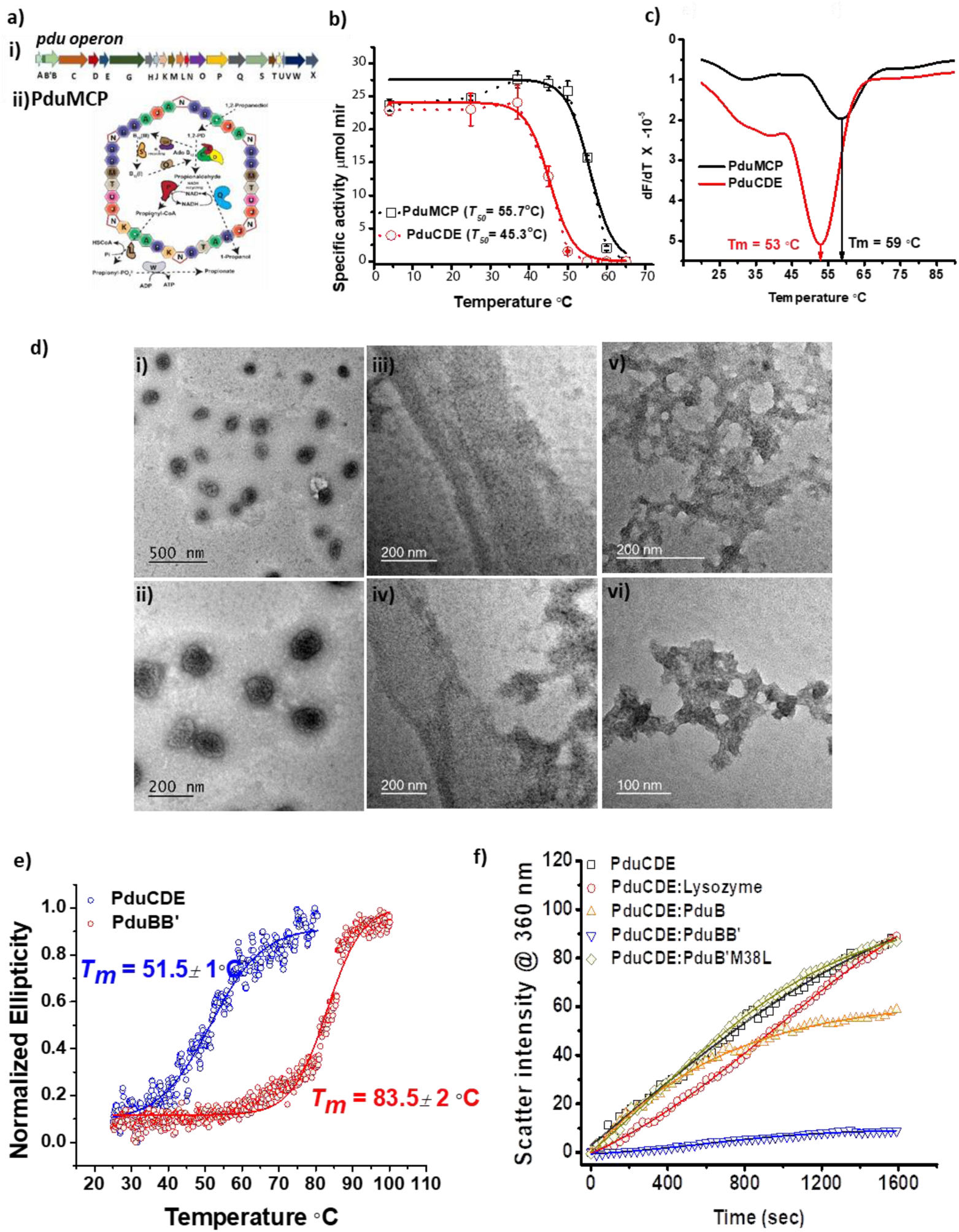
(a) Schematic representation of *Pdu operon* (i) and PduMCP (ii), (b) Temperature dependant diol dehydratase assay using PduMCP (black) and PduCDE (red), (c) Derivative plot of change in intrinsic fluorescence as a function of temperature for PduMCP (black) and PduCDE (red), (d) TEM image of PduMCP (i) and (ii), PduBB’ (iii) and (iv), PduCDE(v) and (vi), e) CD melt of PduBB’ (red) and PduCDE (blue), (f) Aggregation kinetics of PduCDE alone (black), PduCDE in the presence of PduBB’ (blue), PduB (yellow), PduB’M38L (green) and lysozyme (red) at 45°C.

As these MCPs are made up of thousands of protein subunits, understanding the structural and functional roles of each would tell the ways they provide selective advantage to the bacteria. Available reports suggest a vital role of the outer envelope in encapsulating the enzyme system and serving as a semipermeable membrane between the cytosol and internal lumen (12). In the case of PduMCP, the substrate 1,2-propanediol enters the MCP from the cytosol of bacteria through the pore of the shell protein PduA (13), and is further metabolized via a toxic volatile intermediate propionaldehyde. A PduMCP related diol-dehydratase enzyme, PduCDE catalyzes the conversion of 1,2-propanediol into propionaldehyde, which is further utilized by downstream pathways **(Fig. 1a)**. The intermediate product, propionaldehyde is volatile and cytotoxic and may destroy the bacteria if it escapes out of MCP (6). Simulation studies have shown that propionaldehyde experiences a higher free energy barrier as compared to 1,2-propanediol during its movement through the pore of PduA, which blocks its escape out of the MCP (14). Thus, encapsulation limits the escape of toxic intermediate out into the cytosol, protecting the bacteria against toxicity. While this information throws a light on the functioning of one of the shell proteins, it also encourages us to study the properties and role of other shell proteins in the PduMCP.

PduBB’is the most abundant shell protein and makes up 50% of the Pdu-MCP outer shell. It is encoded by the second gene in the *pdu* operon (15) **(Fig. 1a).** It comprises of two proteins PduB’ and PduB that are encoded by two overlapping genes (7). PduB’ (270 amino acids, 28 kDa) is longer than PduB (233 amino acids, 25 kDa) having an extended 37 amino acids long N-terminal region **(Fig. S1a)**. This extended N-terminal region has been suggested to bind the outer shell of PduMCP with its encapsulated enzymes (16). Interestingly, gene deletion studies show that deletion of the PduB/B’ gene from the *pdu* operon, results in the complete abolition of the PduMCP formation (17). While there are reports on the importance of terminal regions of Pdu proteins in the assembly of PduMCP (16,18-20), the impact of the shell proteins on the catalytic efficiency and stability of the enzymes in a confined environment of MCP is not understood. It is important to concede that the formation of complex structures such as MCPs would indeed require interactions between the shell proteins and the enzymes. We hypothesize that, due to limitation in volume (a few femto-liters for a folded MCP), the enzymes would be surface confined onto the shell proteins and this would influence their catalytic efficiency and stability. In this context, we explore the effect of the major shell protein PduBB’ on the stability and activity of the signature enzyme diol dehydratase PduCDE. Our work delineates the importance of PduB and PduB’ combination in the context of PduMCP assembly and enzyme stability, and propose a new role of shell protein PduBB’.

## Results

### Encapsulation provides thermal stability to enzymes in PduMCP

We begin our study by questioning the impact of encapsulation on the stability of enzymes within PduMCP. To address this, we perform temperature dependent enzyme assays using PduMCP and its signature enzyme diol-dehydratase PduCDE in the range 4-50°C. We observe a rise in the specific activity for the PduMCP till 37°C, in contrast to the free enzyme which remains constant (**Fig 1(b), Table S1)**. The enhancement in the specific activity for the PduMCP from 4-37°C is close to 15%. This enhanced activity is retained till 50°C after which there is a gradual drop. The trend is indicated in **Fig. 1b** by dotted lines. Post 60°C the PduMCPs do not show any activity. In contrast, the free enzyme starts to lose its activity immediately after 37°C.

As we observe significant difference in the temperature dependent enzyme activity of the bare and the encapsulated enzyme, we next test the thermal stability of the two systems using intrinsic tryptophan fluorescence of the component proteins as a probe. In case of PduMCP, the shell proteins are deficient in tryptophan residues **(Table S2)**. On the other hand, the Pdu enzymes are rich in tryptophan residues (5 tryptophan in PduCDE). Therefore, Pdu enzymes would contribute maximum to the intrinsic fluorescence of PduMCP and help us to assess the conformational stability of the encapsulated enzymes at higher temperatures. **Fig 1 (c**) shows the derivative plot of fluorescence at λ_max_ with respect to temperature for PduMCP and PduCDE. The Tm of PduMCP (59°C) is higher than that of PduCDE (Tm = 53°C). Also, the Tm values of both PduMCP and PduCDE are around temperatures at which their respective specific activities are completely compromised. This shows that the enzyme PduCDE has a higher conformational stability when encapsulated within PduMCP than in free form. The free PduCDE tend to unfold at lower temperatures and undergoes denaturation. These observations suggest that encapsulation indeed provides thermal stability to enzymes in PduMCP. This adds a new virtue of compartmentalization paradigm in bacteria to the existing list of segregation of metabolic pathway and isolation of toxic intermediates. These results motivate us to dig out the reason behind such enhanced thermal stability of enzyme within PduMCP.

### Shell protein PduBB’ shows chaperone like behavior towards PduCDE

In this section, we take into consideration, the interactions between the outer shell and core enzymes in PduMCP, and ask if shell proteins contribute to enhanced thermal stability of enzymes. We address this query by studying the role of shell protein in providing stability to the enzymes in PduMCP with special emphasis to the shell protein PduBB’. The purified PduMCPs form closed compartments of the size in the range of 100-150 nm (**Fig. 1d (i),(ii)**). PduBB’ comprises half of the outer shell of PduMCP and forms extended sheet when overexpressed and purified **(Fig. 1d(iiii),(iv)).** The diol dehydratase enzyme, PduCDE is a globular protein and purified in three subunits post overexpression in *E. coli* system **(Fig. 1 d (v),(vi) and S1a)**. Our initial characterization show that both the proteins are folded (**Fig. S1b**) and the melting temperature of PduBB’ is ∼80°C and PduCDE is ∼50°C **(Fig. 1e)**. A high melting temperature of PduBB’ shows its ability to resist thermal denaturation. We wonder if this shell protein could also protect Pdu enzyme under thermal stress.

We perform Rayleigh scattering experiment to study the time dependent aggregation of PduCDE in the absence and presence of the shell protein PduBB’. **Fig. 1f** shows a plot of Rayleigh scatter intensity versus time for the PduCDE in the absence and presence of PduBB’ at 45°C. In the absence of the shell protein, the scatter intensity increases with time indicating aggregation of PduCDE due to its thermal denaturation. Interestingly, we do not observe such enhancement in the scatter intensity in the presence of the shell protein PduBB’, which suggests that the shell protein PduBB’ curbs the unfolding of the enzyme PduCDE, preventing its thermal denaturation. Since PduBB’ is a combination of PduB and PduB’, we question their individual role in providing thermal stability to the enzyme. To get PduB’ alone, we generate a single mutant clone PduB’M38L **(Fig. S1a).** Both PduB and PduB’M38L fail to prevent the aggregation of PduCDE, highlighting the significance of this combination in PduMCP. To validate our observation, we repeat our experiment using a non-native protein, lysozyme which is not known to provide thermal stability to biomolecules in living system. As shown in **Fig. 1g**, PduCDE tends to aggregate in the presence of a non-native protein lysozyme, eliminating the role of non-specific interactions in our observations. This experiment shows that PduBB’ has a unique property to prevent thermal denaturation of the native enzyme PduCDE.

The high thermal stability of the shell protein PduBB’ and its chaperone like activity intrigues us to look for the evolutionary origin of such a thermostable shell protein in a mesophilic *Salmonella enterica*. Our bioinformatics analysis show that genes for PduMCP appeared in thermophilic firmicutes and were horizontally transferred to the proteobacterial species. The thermostable property of the shell protein PduBB’ has been passed down to *Salmonella enterica* by its thermophilic ancestors (**Supporting Information and Fig. S2**).

### PduCDE shows enhanced activity in the presence of the shell protein PduBB’

The next question we asked is, besides providing thermal stability to PduCDE, does PduBB’ also influence its catalytic activity. We look for the activity of PduCDE in the absence and presence of PduBB’. At lower PduBB’ to PduCDE ratio (w/w) we do not observe any significant change in the enzyme activity, however, upon increasing the ratio to more than 8:1, a significant change is noticed **(Fig. 2a)**. This motivates us to check if PduBB’ also influences the activity of PduCDE post thermal stress at higher temperatures. Interestingly, post heat shock at 45°C, PduCDE shows an activity (23.73±0.79 µmol min^-1^ mg^-1^) in the presence of PduBB’ but a significantly lower activity (13.29±1.91 µmol min^-1^ mg^-1^) in its absence (p ≤ 0.05) **(Fig. 2b)**. While we do not observe any PduCDE activity post heat shock at 50°C, the enzyme shows an activity of 4.24±0.63 µmol min^-1^ mg^-1^ in the presence of PduBB’ **(Fig. 2b)**. This experiment shows that the shell protein PduBB’, not only increases the catalytic activity of PduCDE under optimum conditions but also protects the enzyme during thermal stress. However, presence of a non-native protein lysozyme results in a decreased catalytic activity **(Fig. S3a).** Decreased activity can be a result of decrease in the accessibility of the substrate to the enzyme due to the presence of non-native protein in its vicinity. This observation again eliminates the role of non-specific interactions in our observations. We also find that the shell protein PduBB’ fail to influence the catalytic activity or provide stability to a non-native enzyme Alcohol dehydrogenase **(Fig. S3b)**. The constituent shell proteins of PduBB’, PduB and PduB’M38L when tested alone for the chaperone like activity, fail to influence the catalytic activity of PduCDE under optimum condition **(Fig. 2c)** and post thermal shock **(Fig. 2d)**. This suggests that the presence of both PduB and PduB’ is crucial for the protective role of PduBB’. To unearth the underlying reason behind this observation, we need to understand how different is PduBB’ from its constituent proteins, in terms of assembly, stability and affinity towards PduCDE.

**Fig 2:**
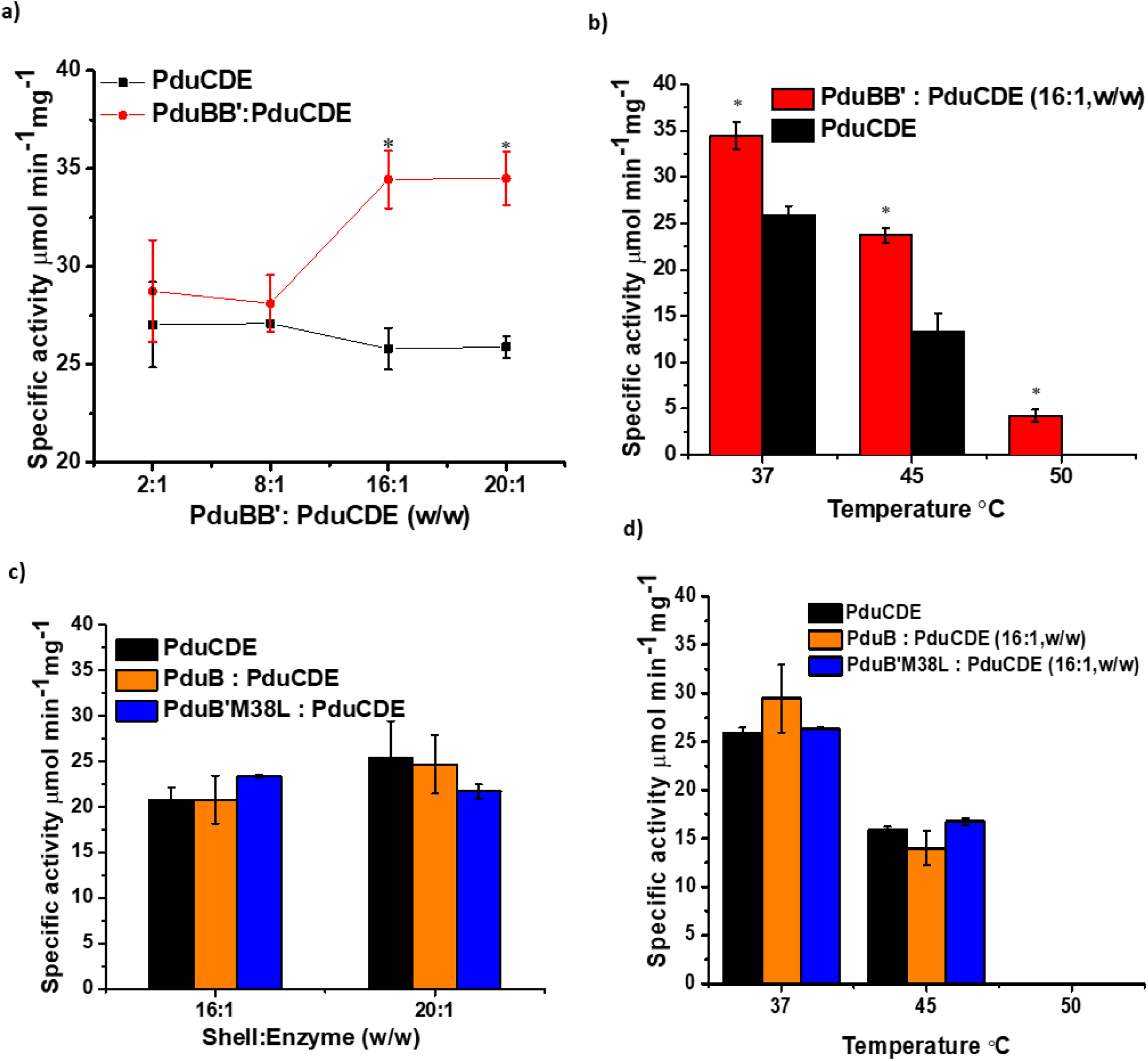
(a) Specific activity of PduCDE in the absence (in black) and presence (in red) of PduBB’ at different w/w ratio, (b) Specific activity of PduCDE in the absence (in black) and presence (in red) of PduBB’ at PduBB’: PduCDE ratio of 16:1 (w/w) post heat shock at 37°C, 45°C and 50°C, (c). Specific activity of PduCDE in the absence (in black) and presence of PduB (in orange) and PduB’M38L (in blue) at shell:enzyme ratio of 16:1 and 20:1, (d) Specific activity of PduCDE in the absence (in black) and presence of PduB (in orange) and PduB’M38L at shell:enzyme ratio of 16:1 (w/w) post heat shock at 37°C, 45°C and 50°C [* p ≤ 0.05].

First, we explore the differences between the three variants of PduB* in terms of their assembly and stability. An important difference between PduBB’ and PduB, is the presence of the extended 37 amino acids N-terminal in PduB’ **(Fig. S4)**. We observe that PduB has a tendency to form soluble associates *in vitro*, which separates out of solution as condensate indicated by a high optical density of PduB at 600 nm (**Fig. 3a)**. Both PduBB’ and PduB’M38L, with 37 amino acids N –terminal, remain soluble and do not form any condensate. Interestingly, the turbidity observed for PduB disappear upon heating at 60°C for 10 minutes **(Fig. 3a)** while the secondary structure of the protein obtained from CD spectra are similar indicating that heating does not alter the protein folding **(Fig. 3b).** Lower ellipticity of PduB prior to heating is due to the formation of self-associates of protein (21). These observations reveal that the 37 amino acids N-terminal region has a vital role in providing solubility to the shell protein PduBB’. In the absence of the N-terminal region, PduB fails to remain soluble and form large associates in solution. A comparison of the secondary structure of PduBB’ and PduB’M38L at same concentration using circular dichroism, shows lower ellipticity for PduB’M38L **(Fig. 3c)**. Next, we compare the thermal stability of PduB and PduB’M38L with that of PduBB’. We monitor the secondary structural changes of the three forms of PduB with temperature. PduBB’ retains its secondary structures up to 80°C **(Fig. 3d).** PduB also shows high thermal stability, retaining its secondary structures up to 80°C **(Fig. 3e)**. Both PduBB’ and PduB show a sharp decrease in ellipticity measured at 222 nm above 80°C **(Fig S5a and S5b)**. PduB’M38L shows a gradual decrease in ellipticity with the increase in temperature **(Fig. 3f)**. A plot of ellipticity measured at 222 nm versus temperature shows a prominent transition around 60°C, followed by a further decrease in ellipticity **(Fig. S5c)**, indicating lower stability of the shell protein. Together our results suggest that the presence of both PduB and PduB’ is necessary for the formation of a stable and soluble shell protein PduBB’. The difference in the solubility and stability of the PduB and PduB’M38L leads us to hypothesize that the absence of either of these two affects the self-assembly property of PduBB’. We test this hypothesis by studying the self-assembly behavior of the three variants of PduB* using dynamic light scattering (DLS) as discussed in the following section.

**Fig 3:**
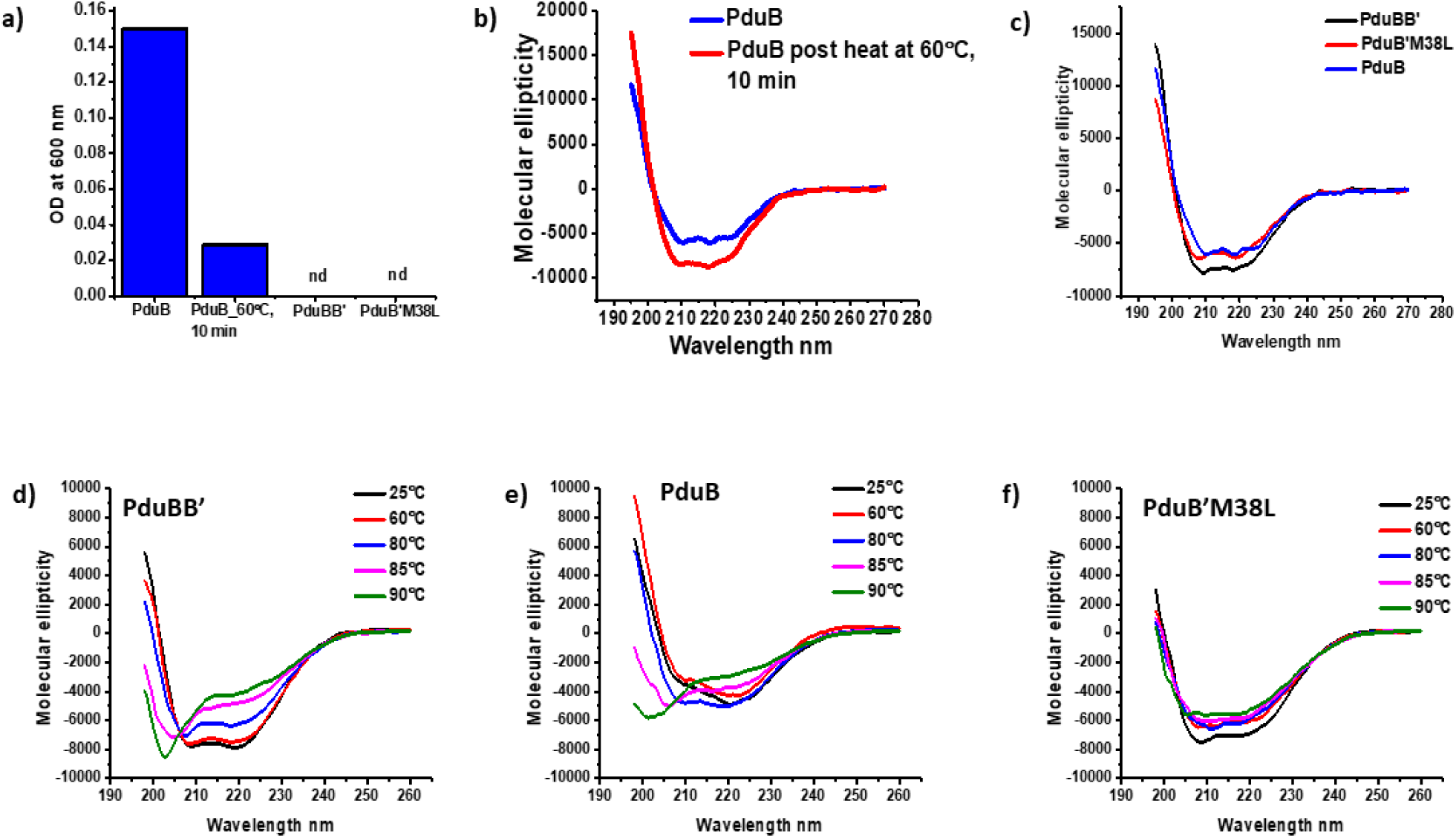
(a) Scattering at 600 nm for PduB, PduBB’ and PduB’M38L (0.25 mg/ml each), pre and post heating at 60°C and the CD spectra of PduB (0.25mg/ml) pre and post heating (inset), (b) CD spectra of PduBB’ (0.25 mg/ml) and PduB’M38L (0.25 mg/ml) and PduB (0.25 mg/ml (b), CD melt spectra of 0.25 mg/ml of (c) PduBB’, (d) PduB and (e) PduB’M38L

### PduBB’ balances the self-assembly properties of PduB and PduB’

In this section our main focus is to understand the difference in the self-assembly behavior of the three variants of PduB shell proteins in solution phase. **Fig. S6** shows the size distribution of the three shell protein at different concentrations. PduB forms large associates even at lower concentration of 0.1 mg/ml **(Fig. S6a)** and has a maximum population of associates in micron range **(Fig. S6b)**. Increase in concentration has no significant effect on the size distribution of PudB **(Fig. S6a)**. PduB has a high tendency to self-associate and do not form a soluble and stable shell assembly. PduB’M38L shows size distribution between 100 to 300 nm **(Fig. S6c)** and has a maximum population of shell assemblies of size less than 20 nm **(Fig. S6d)**. We do not see any size distribution in the micron range at higher concentrations **(Fig. S6c).** This suggests that PduB’M38L has a very less tendency to self-associate. We make interesting observations in case of PduBB’. An increase in the shell protein concentration produce size distributions both around 100 nm and in the micron range **(Fig. S6e)**. At a concentration of 0.8 mg/ml, around 50 % of the size distribution is in the micron range and rest is around or less than 100 nm **(Fig. S6e)**. PduBB’ also has a maximum population of shell assemblies less than 100 nm **(Fig. S6f)** but unlike PduB’M38L, we do observe size above 1000 nm **(Fig. S6e)**. This suggests that PduBB’ balances the self-assembly properties of both PduB and PduB’. Concentration dependent CD spectra shows that all the three shell proteins retain their secondary structures at different concentrations **(Fig. S7)**. The overall ellipticity of PduB and PduB’M38L is less than that of PduBB’, supporting our earlier results **(Fig 3c).** Interestingly, the ellipticity of PduBB’ decreases with the increase in shell protein concentration **(Fig. S7a).** We attribute this decrease in ellipticity to the formation of higher order assemblies at higher concentrations, as demonstrated by DLS results **(Fig S6e).**

A naturally optimized, stable self-assembly in case of PduBB’ would provide the suitable scaffold necessary for enhancing the activity and stability of the enzyme PduCDE. The shell protein PduB is N-terminal truncated and undergoes self-association, while PduB’M38L with the N-terminal region has the least tendency to self-associate. An auto-optimized self-association in case of PduBB’, suggests a vital role of N-terminal region in regulating the self-assembly of PduBB’ post translation which perhaps has implications in outfitting the size of the mature PduMCP.

### PduCDE has a greater affinity towards PduBB’ than PduB or PduB’M38L

The three shell proteins PduBB’, PduB and PduB’M38L differ in their assembly and stability as suggested by our CD and DLS results. Now, we probe the interactions between the enzyme and the shell proteins using biolayer interferometry. We immobilize the enzyme on to a sensor tip (see experimental procedures for details) and monitor its interactions with the shell proteins present in free solution. **Fig 4** shows association and dissociation kinetics between the shell proteins PduBB’/B/B’M38L and the enzyme PduCDE.

**Fig 4:**
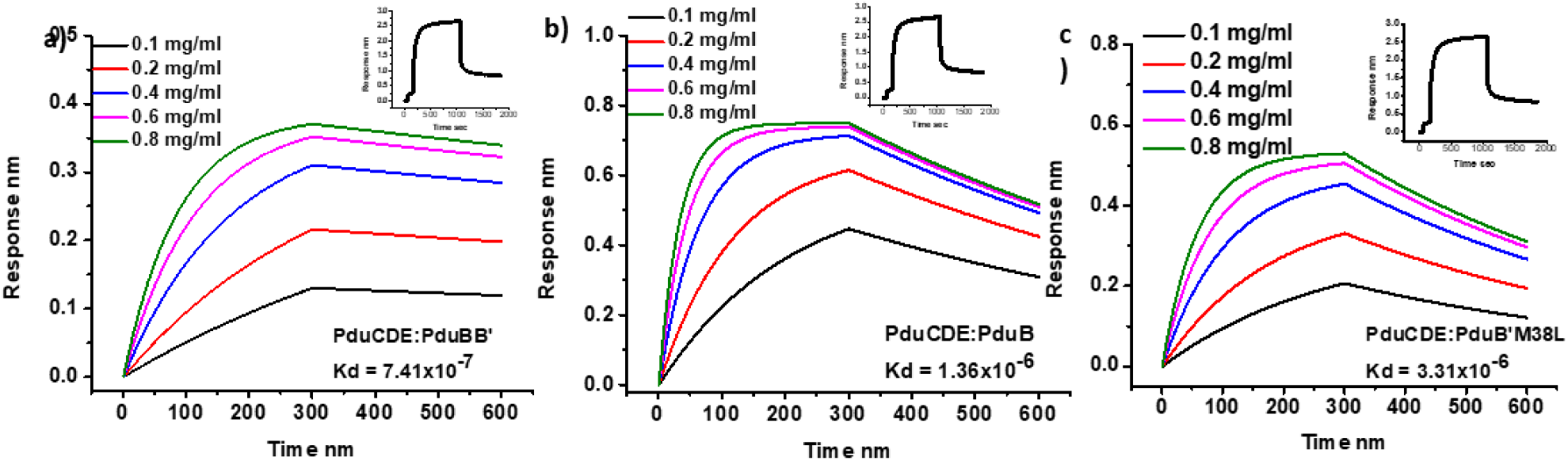
Interactions between PduCDE and the shell proteins (a) PduBB’, (b) PduB and (c) PduB’M38L. PduCDE is immobilized using 1-ethyl-3-(3-dimethylaminopropyl) carbodiimide (EDC)/N-hydroxysuccinimide (NHS) method onto the amine reactive second generation (AR2G) biosensors and its association with increasing concentrations of shell proteins is probed. The loading efficiency of PduCDE on to the sensor is shown in inset.

In case of PduBB’ **(Fig 4a)**, the association response increases with the increase in the concentration of the shell protein. A moderate affinity exists between the shell proteins and the enzyme with dissociation constant in micromolar range). This explains the reason behind the need for high shell/enzyme ratio for the shell protein to enhance the activity and stability of the enzyme PduCDE. In case of PduB **(Fig 4b)**, we see a greater association response compared to PduBB’. This is because of the larger associates of PduB that interact with the enzyme immobilized sensors. However, with the increase in PduB concentration, the association response tends to saturate, followed by faster decay kinetics. This indicates non-specific and weaker interactions between the enzyme and PduB at higher concentrations. The association response of PduB’M38L is comparable to that of PduBB’ as they are both soluble in aqueous solution. Like PduB, it also shows faster decay kinetics at higher concentrations indicating weak interactions between PduB’M38L and PduCDE. **(Fig 4c)**. Global fitting of the association and dissociation kinetics reveals Kd values for PduCDE:PduBB’ PduCDE:PduB and PduCDE:PduB’M38L to be to be 7.41*10^−7^, 1.36*10^-6^ and 3.31*10^-6^ respectively. This suggests that PduCDE has a stronger affinity towards PduBB’ than its individual components (PduB and PduB’M38L). We perform a control experiment using BSA and PduCDE **(Fig S8a)** and observe a saturated association response with the increase in BSA concentration, suggesting non-specific interactions between PduCDE and BSA. We also ensure that the His-tag associated with our proteins do not interfere in our binding experiment. To do this, we use a His-tagged non-native protein Dmrb and monitor its association with PduCDE immobilized sensor. No association response is recorded between Dmrb and PduCDE **(Fig S8b)**, eliminating the role of His-tag in our experiment. In the following section, we probe the interaction between PduCDE and three variants of PduB*, with both the enzyme and the shell protein in solution phase.

### Restricted motion of Shell proteins in the presence of enzyme PduCDE

In this section, we study the change in the dynamics of shell protein in the vicinity of the enzyme in solution. Fluorescence anisotropy is a very useful technique which not only help us to probe protein-protein interactions but also gives us information regarding the changes in structural dynamics of a protein during interactions (22).

Due to extended sheet formation exhibiting high resilience, we expect a slow global motion in case of shell protein. PduBB’ contains two cysteine residues, Cys^43^ and Cys^187^. Labeling these cysteine residues with a fluorophore must allow us to probe the rotational motion of the shell protein. Since the crystal structure of PduBB’ from Salmonella enterica is not solved, we perform homology modeling using Swiss model server (23,24). While the model clearly indicates the position of Cys^187^ in the beta strand, it eliminates the extended N-terminal region including Cys^43^ **(Fig. S9 a)**. The extended N-terminal region was eliminated possibly due to the absence of the extended N-terminal region in PduB of *Lactobacillus reuteri*. We perform *ab initio* modelling of PduB’ and PduB using AIDA server (25).

Interestingly, both the models show Cys^43^ to be a part of a coil in the N-terminal and Cys^187^ to be a part of beta strand **(Fig. S9 b)**. Since, beta strand is more rigid, we expect Cys^43^ in the coiled region to contribute maximum to the anisotropy of the shell protein. The shell protein PduBB’ is labeled with a thiol reactive fluorophore Acrylodan (26) and time resolved anisotropy for the shell protein is carried out in the absence and presence of the enzyme PduCDE. The shell protein PduBB’ is mixed with PduCDE at equimolar ratio prior to the anisotropy measurements. **Fig. 5a** shows the time resolved anisotropy for the dye labeled PduBB’ in the presence and absence of the enzyme. The decay kinetics indicate a restricted motion of the shell protein in the presence of PduCDE. The decay curves are best fitted with two exponential function (*Eq. 1*) and two sets of rotational co-relation time are obtained. The correlation time along with their corresponding amplitudes are shown in **Table S3**. In the absence of PduCDE, we get one correlation time in sub-nano second range (0.58 ± 0.43 ns for PduBB’ and 0.35 ± 0.82 ns for PduBB’:PduCDE) corresponding to the motion of the acrylodan molecule **(see Fig. S9c)** bound to the protein. The second correlation time is longer (5.19 ± 2.50 ns for PduBB’ and 10.69 ± 1.03 ns for PduBB’+PduCDE)). In order to understand the origin of the motions contributing to these longer co-relation time, the molecular weight of the protein needs to be taken into consideration. Jain et al. 2016 (27) reported a rotational correlation time for the global motion of beta-microglobulin (MW ∼ 12 kDa) around 5 ns. In another study, Zhou et al. 2012 (28) attributed a longer correlation time (above 30 ns) to the regional dynamics of Troponin I (MW ∼ 23 kDa). As mentioned earlier, PduBB is a trimer of PduB’ (28 kDa) and PduB (25 kDa) and exists as an extended sheet. The sheets must have a slow rotational correlation time. Also, the average life time of acrylodan is 4 ns, hence, negligible contribution will be made by the tumbling motion of large sheets to the anisotropy during the time fluorophore is in excited state. Since Cys^187^ is a part of a rigid beta strand, we attribute the correlation time around 5 ns to the segmental motion of the coil region (Cys^43^) in the N-terminal of PduBB’. Increment in the co-relation time to ∼10 ns indicates restricted flexibility of this region.

**Fig 5:**
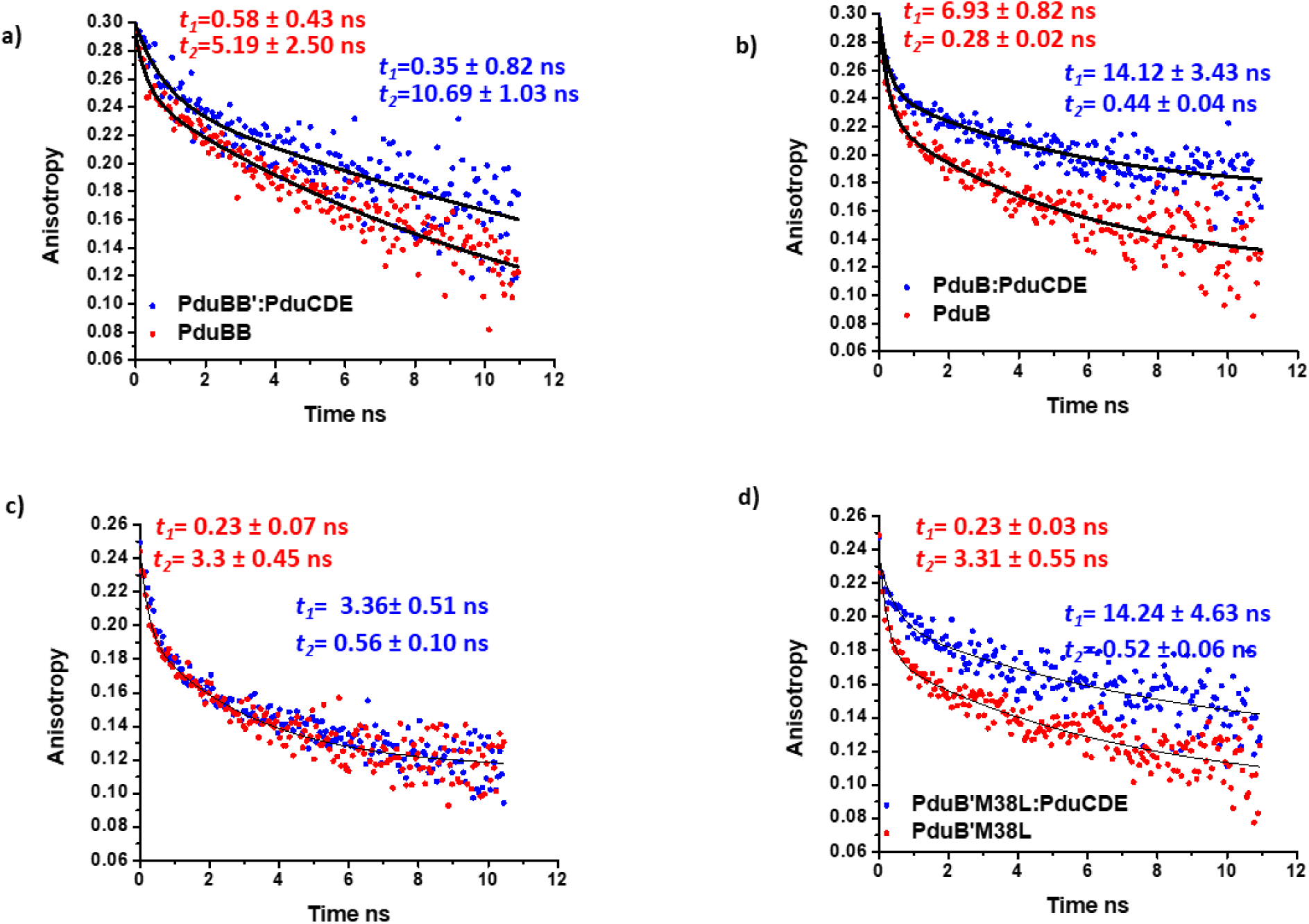
(a) Time resolved anisotropy decay of shell protein PduBB’ in the absence and presence of enzyme mixed in 1:1 molar ratio, (b) Time resolved anisotropy decay of shell protein PduB in the absence and presence of enzyme mixed in 1:1 molar ratio. (c) Time resolved anisotropy decay of PduB’M38L in the absence and presence of the enzyme PduCDE mixed in 1:1 molar ratio and (d) Time resolved anisotropy decay of PduB’M38L in the absence and presence of the enzyme PduCDE mixed in enzyme: shell of 4:1 molar ratio.

We get similar result in case of PduB **(Fig. 6b)**, a faster correlation time (0.28 ± 0.02 ns in case of PduB and 0.44 ± 0.04 ns in case of PduB:PduCDE), corresponding to the fluorophore tumbling and a longer correlation time (6.93 ± 0.82 ns in case of PduB and 14.12 ± 3.43 ns in case of PduB:PduCDE). Since PduB forms large associates in aqueous medium, the longer correlation time around 6 ns must arise from the coil region in the N-terminal end of PduB. A restricted motion of this region in the presence of PduCDE is evident from the increment in the longer correlation time to ∼14 ns. Our DLS results reveal that PduB’M38L has the least tendency to self-associate **(Fig. 5b)**. Therefore, we expect the N-terminal region in case of PduB’M38L to be least restricted exhibiting a faster tumbling motion. Interestingly, we do not see any difference in the decay kinetics of PduB’M38L in the absence and presence of PduCDE in equimolar ratio, indicating weaker interaction between the two **(Fig. 5c)**. We repeat the experiment by increasing the concentration of PduCDE **(Fig. 5d)**. The longer correlation time in case of PduB’M38L is ∼ 3 ns corresponding to the flexible coiled region near the N-terminal end of the shell protein. In the presence of PduCDE (shell:enzyme, 4:1 molar ratio), the longer correlation time increases to ∼14 ns, suggesting reduced flexibility near the N-terminal region of the shell protein. Together our anisotropy results show a restricted motion of the shell proteins PduBB’, PduB and PduB’M38L in the presence of the enzyme PduCDE, confirming the interactions between the shell proteins and the enzyme. These results suggest that all the three shell proteins PduBB’, PduB and PduB’M38L can bind to the enzyme PduCDE. However, it’s PduBB’ which has a higher affinity towards PduCDE, as shown by BLI result **(Fig 4a)**. Also, due to a well-balanced self-assembly **(Fig. S6e)**, the combination of both PduB and PduB’ in PduBB’ would provide a suitable scaffolding necessary for the improved catalytic activity and enhanced thermal stability of the enzyme.

**Fig 6:**
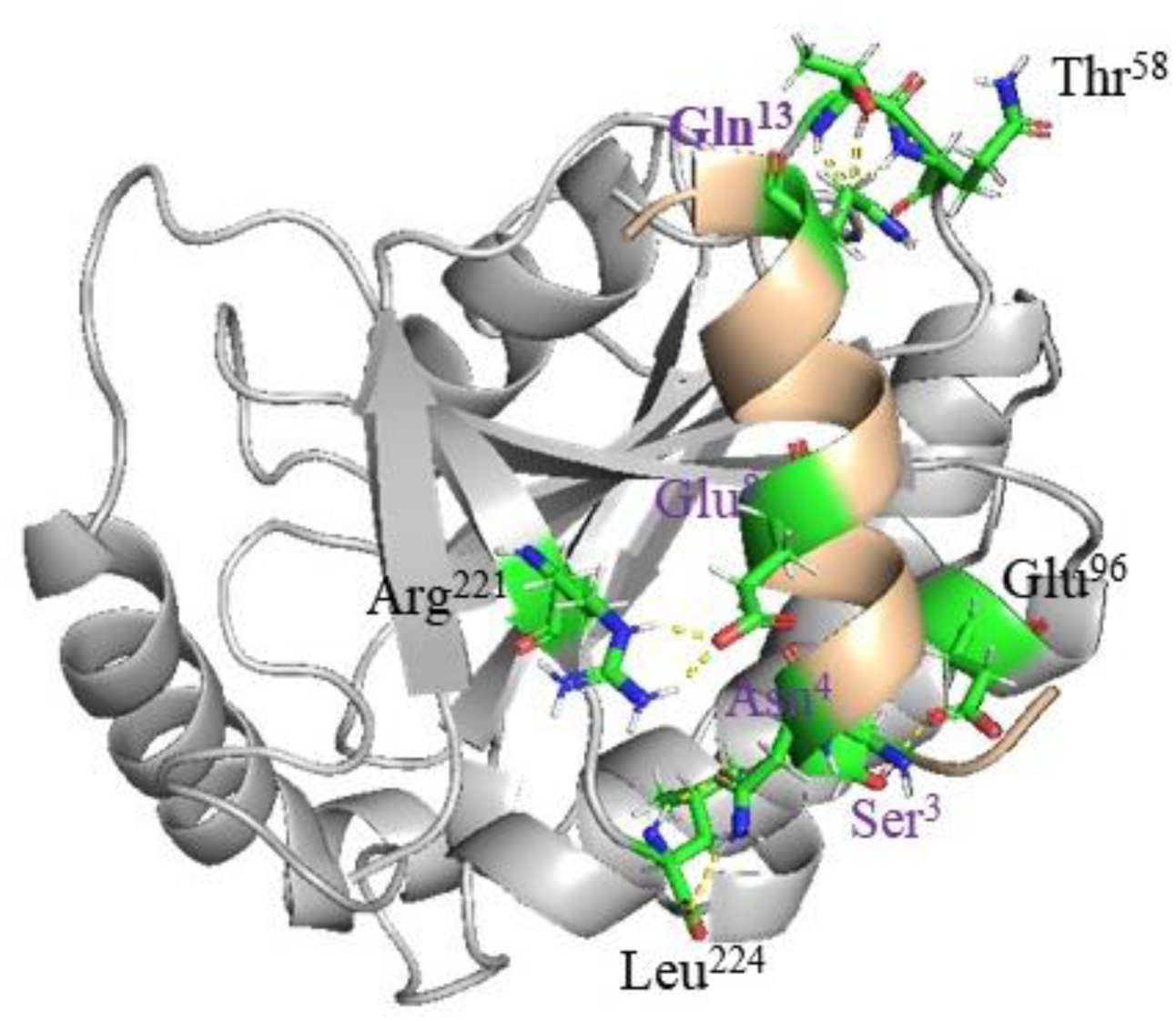
Peptide-protein docking between first 15 amino acids of N-terminal of PduB’ with D subunit of diol dehydratase (1DIO), using PIPER-Flex Pep Dock. The residues Ser3, Asn4, Glu8 and Gln13 in the N-terminal of PduB’ (in purple) form hydrogen bonds with Glu96, Leu224, Arg221 and Thr58 (in black) respectively of subunit D of the enzyme diol dehydratase. A salt bridge also exists between Glu8 of PduB’ and Arg221 of subunit D.

### Predicting residue level interactions between PduBB’ and PduCDE

The anisotropy results help us hypothesize the presence of a functional region near the N-terminal of the shell protein PduBB’ that participates in binding with the enzyme PduCDE. It is also important to understand how the absence of the extended 37 amino acids N-terminal region in PduB alters the binding sites. Due to the absence of solved crystal structure of PduBB’, we go for homology modeling of the shell protein followed by molecular docking with the enzyme. This is done to enhance our understanding about the shell-enzyme interactions observed in our study. We perform molecular docking of Diol-dehydratase (PDB Id: 1DIO) with the modeled shell proteins PduB and PduB’ using ClusPro docking server (29-31). The extended N-terminal region of PduB’ binds to the subunit D of the enzyme **(Fig. S10a)**. 10 hydrogen bonds (**Table S4**) and 1 salt bridge hold the complex together. We use PIPER-Flex PEP dock (32-34) to see the interactions between first 15 amino acids of N-terminal region of PduB’ with D subunit of the enzyme **(Fig. 6)**. The N-terminal region is automatically folded into a helix and bind to subunit D through hydrogen bonds and a salt bridge. The hydrophobic residues like V7 and I10 are completely buried (**Table S5**), suggesting the role of hydrophobic interactions in mediating shell enzyme interactions. Binding of the N-terminal region to the enzyme would restrict motion of this region. This explains the increment in the longer rotational co-relation time seen in case of PduBB’ and PduB’ in the presence of the enzyme PduCDE.

In the absence of the 37 aa N-termainal region, the binding sites between the shell protein and the enzyme are altered **(Fig. S9b)**. 12 hydrogen bonds (**Table S6**) and 2 salt bridges hold the complex together. The residues E3, K4 and S5 in the coil region of the N-terminal of PduB participates in hydrogen bonding with the subuit D of the enzyme. E3 also forms a salt bridge with K199 of subunit D. These interactions would restrict the rotational motion of this coiled region increasng the rotational co-relation time in the presence of the enzyme, seen in our anisotropy experiment.

It is noteworthy that the crystal structure of the diol-dehydratase enzyme which has been used in the docking studies, lacks the extended N-terminal region. The N-terminal region of PduD is necessary for its encapsulation within PduMCP (19). We dock first 15 amino acids of PduD N-terminal against modeled PduB trimer using PIPER-Flex PEP dock. The N-terminal of PduD is automatically modeled into a helix that goes and bind in the pore region on the concave side of PduB **(Fig. S11a and S12a).** The hydrophobic residues face the cleft of the pore and charged amino acids face outside and are exposed to the solvent. This is evident from very low solvent accessible surface area for hydrophobic residues (**Table S7**). The orientation of hydrophobic residues towards the cleft of the pore suggests the role of these residues in binding to the concave face of PduB shell protein. Interestingly, when docked onto the convex surface of PduB trimer **(Fig. S11b and S12b)**, the binding site of the N-terminal of PduD is slightly towards the edge of the trimer. Also, there is an active involvement of charged amino acids in the formation of hydrogen bond with the shell protein (**Fig. S12b**), suggesting a role of electrostatic interactions in the interaction of N-terminal of PduD with the convex surface of PduB.

## Discussion

PduMCPs are very small in size (100-150 nm) in which enzymes are restricted to a very limited space and volume (9). The interactions between the outer shell and the enzymes are essential to the formation an intact MCP (18-20). Our work elucidates the link between molecular confinement and conformational stability of enzymes in PduMCP. In free solution PduCDE is very susceptible to heat and lose its activity by more than 40% at 45°C. While in encapsulated PduMCP, the activity of the enzyme falls by more than 40% at 55°C. This shows that the enzymes tend to be thermally more stable in PduMCP than in free aqueous environment. In this regard, the role of shell protein PduBB’ is investigated. We provide evidence in support of the idea that PduBB’ has the ability to improve the catalytic activity of PduCDE and provide protection to the enzyme under thermal stress. Although we do not see any effect of PduBB’ on PduCDE activity at lower shell:enzyme ratio, around 33% enhancement in the activity is seen at higher shell:enzyme ratio. This moderate enhancement can be explained by the fact that **Fig 2(a)** represents a free ensemble system and not an encapsulated system. Various factors such as substrate channeling or substrate accessibility to the enzyme would be better in an encapsulated system than in a free ensemble system. Besides, BLI results also suggest a moderate affinity between the shell protein and enzyme. However, post thermal shock at 45°C, the PduCDE activity in the presence of PduBB’ is around 78% higher than PduCDE in the absence of the shell protein, suggesting the protective role of the shell protein. Aggregation kinetics experiment confirms that PduBB’ prevents the unfolding and aggregation of the enzyme PduCDE. We also perform these experiments using non-native proteins lysozyme to ensure that these results are not due to non-specific interactions between the shell protein and the enzyme. The shell protein PduBB’ has a tendency to resist thermal denaturation. Due to a high similarity in sequence of PduB from different enterobacterial species **(Fig. S13**), we propose that this thermo-tolerant property of the shell protein must have been preserved in other enterobacterial species.

Among all the shell proteins in PduMCP, PduBB’ is the only one which is the product of two overlapping genes. PduBB’ is a trimer and is a combination of PduB and PduB’. Through our work we show that the simultaneous expression and assembly of the PduB and PduB’ is highly important for the formation of a stable and soluble shell protein, and that this combination has a positive influence on the efficiency and stability of the signature enzyme of PduMCP. Interaction studies also confirm that PduCDE has a stronger affinity towards PduBB’ than its individual component shell proteins. Lower solubility of 37 amino acids N-terminal truncated PduB suggests that, beside participating in binding shell to the enzyme core in PduMCP (16), the N-terminal region also contributes to the solubility and self-assembly of PduBB’. Interestingly, the N-terminal region of middle and smaller subunits of diol dehydratase lowers the solubility of the enzyme (35). Identifying the binding shell partners for the enzymes is the key question in MCP research for the development of protein based hybrid materials. Our study shows that the major shell protein PduBB’ acts as scaffold for PduCDE, enhancing its activity and stability. This observation is indeed encouraging and may lead to the development of MCP based 2-D and 3-D thermostable nano-materials using genetic engineering and chemical biology approaches. These all-protein smart biomaterials can be an alternative to the confinement of enzymes via immobilization onto artificial matrices like silica (36), glyoxyl-agaraose (37) or epoxy-activated supports (38). A question that still remains regarding the PduBB’ is that whether the combination is a heterotrimer or a dimer of trimers of PduB and PduB’. Recently, Sommer et al 2019 have shown that in beta-carboxysomes, shell proteins Ccmk3 and Ccmk4 exist as heterotrohexamers and alter the permeability properties of carboxysome shell (39). This suggests that shell protein combination can have a significant role in the functioning of MCPs. Further insights into the structure of the PduBB’ protein will be needed to understand stoichiometry in which its components assemble.

### Experimental procedures

All the chemicals unless mentioned are procured from Sigma (India). Shell proteins PduB and PduBB’ constructs are gifts from Prof. TA Bobik (Iowa State University, Ames, USA).

#### Purification of PduMCP

We carry out the purification of PduMCP by inducing its expression in *Salmonella enterica* as described earlier (40). 1% of overnight grown culture of *Salmonella enterica* serovar Typhimurium LT2 is inoculated in 400 ml of 1X NCE (non-carbon E) medium, which is supplemented with 0.6 % 1,2 PD, 0.5 % Succinic acid and 1mM of MgSO4 and incubated at 37°C for 16 hours. The harvested cells are washed with Buffer A (50 mM Tris Base pH 8, 500 mM KCl, 25 mM NaCl, 12.5 mM MgCl_2_, 1.5 % 1,2-PD; 8000 X g for 5 min at 4 °C). The washed cells are re-suspended in Buffer A containing 75% bacterial protein extraction reagent (BPER-II), 2 mg Dnase, 0.4 mM phenylmethane sulfonyl fluoride (PMSF) and 1 mg/ml of lysozyme. The re-suspended cells are kept on a shaker at 45-50 rpm at room temperature for 30 min, followed by incubation on ice for 5 min. The lysed cells are removed by centrifugation (12,000 X g for 5 min, 4 °C) and PduMCPs in the supernatant are further pelleted by centrifugation (20000 X g for 20 min, 4 °C). The pellet is re-suspended in Buffer A containing 60% of B-PER II and 0.4 mM PMSF and is centrifuged at 20,000 X g for 20 min at 4 °C. The supernatant is discarded and the thin film obtained is re-suspended in pre-chilled Buffer B (50 mM Tris Base pH 8, 50 mM KCl, 5 mM MgCl2, 1 % 1,2-PD). The re-suspended thin film is centrifuged at 12,000 X g for 5 min at 4 °C. The supernatant containing PduMCP is collected at stored at 4°C.

#### Purification of PduCDE and shell protein

1% of overnight grown primary culture of E.coli BL21DE3 (transformed with PduCDE or shell protein constructs) is inoculated in 400 ml of LB media and incubated at 37°C for 1.5 to 2 hours until it reaches an OD_600_ of 0.5. Protein expression is induced by adding 1 mM IPTG and incubating the culture at 37°C for 4 hours and at 28 °C for 12 hours for enzyme and shell proteins respectively. The cells are harvested and lysed in column buffer (50mM Tris-base pH 7.5, 200mM Nacl and 5 mM imidazole). The supernatant is passed through Ni-NTA column. The column is then washed with washing buffer containing 50 mM imidazole and protein of interest is eluted by passing elution buffer containing 200 mM imidazole. The eluted protein samples are dialyzed in 10 mM sodium phosphate buffer (pH 7.4). Protein concentration is checked using Bradford reagent and purity is checked by performing SDS-PAGE.

#### MBTH diol dehydratase assay

We estimate the diol dehydratase activity of PduMCP and PduCDE by 3-methyl-2-benzothiazoline hydrazine (MBTH) method. 2 μg of PduCDE or 5 μg is added to 900 μl of the assay buffer (0.2 M 1,2-propanediol, 0.05 M KCl, 0.035 M potassium phosphate buffer (pH 8.0) at 37 °C. Reaction is induced by adding 50 μl of adenosyl cobalamin AdoCbl (15 μM) and quenched after 10 min by adding 1 ml of potassium citrate buffer (pH 3.6). 0.5 ml of 0.1 % MBTH is added and the reaction mixture is incubated at 37 °C for 15 min. After 15 minutes 1 ml of double distilled water is added and absorbance of the product formed is taken at 305 nm using UV spectrophotometer. We define specific activity as μmol of product formed by one milligram of enzyme in one minute.

#### Thermal shock assays

To check the effect of temperature on the activity of PduMCP and PduCDE, the proteins samples are exposed to temperatures ranging from 4°C to 65°C for a short duration of 2 min, followed by diol dehydratase assay as described above. To check the effect of shell proteins on the activity of PduCDE enzyme, 1 mg/ml of shell protein is mixed with PduCDE at different w/w ratio and incubated at 4°C for 1 hr. The samples are then subjected to thermal shock at 37°C, 45°C and 50°C for a short duration of 2 min. After cooling down the samples to room temperature, they are used for MBTH diol dehydratase assay.

Alcohol dehydrogenase (Alcdh) assay is carried out at room temperature in 200 µl reaction mixture containing, 150 mM ethanol, 300 µM NAD^+^ and 6µg of enzyme, and production of NADH is monitored by taking absorbance at 340 nm.

#### Fluorescence spectroscopy

Temperature dependent intrinsic fluorescence assay for PduMCP and PduCDE is carried out using Spectrofluorimeter FS5 (Edinburgh Instruments, UK). 0.1 mg/ml of PduMCP or 0.2 mg/ml of PduCDE are excited at 280 nm (band width = 1 nm) and emission spectra (band width = 1 nm) are recoded between 300 to 500 nm.

#### Dynamic light scattering (DLS)

The size distribution of the shell proteins is recorded using a ZetaSizer Nano ZSP (Malvern Instruments, UK). The scattered intensity is measured at backscattered angle of 173°. For each sample, three reading are recorded. The size distribution of the three shell proteins is compared by plotting intensity percentage of the samples at different shell protein concentrations. The maximum population of shell assemblies present in solution of all the three shell proteins is compared by plotting the number percentage of the samples at different concentrations.

#### Circular Dichroism (CD) spectroscopy

CD spectra of shell proteins are recorded using a CD spectrophotometer (Jasco J-1500, CD spectrophotometer, (Jasco, USA). 200 µl of samples are taken in a quartz cell with a path length of (0.1 cm) in a nitrogen atmosphere for measurements in the far-UV region (195-260 nm). An accumulation of three scans with scanning speed of 200 nm/min is performed. Sample temperature is maintained at 25 °C using a mini circulation water bath (Jasco MCB-100) connected to the water-jacketed sample holder chamber. CD spectra are recorded from 25 °C to 100°C at the rate of 2 degrees per minute.

#### Biolayer interferometry

Interactions between the enzyme and shell proteins are carried out using Fortebio OctateK2 (Molecular Devices, USA). Amine reactive second generation (AR2G) biosensors are activated using a solution containing 40mM 1-ethyl-3-(3-dimethylaminopropyl) carbodiimide (EDC) and 20 mM of N-hydroxysuccinimide (NHS) mixed in 1:1 molar ration. This is followed by immobilization of 0.5 mg/ml of PduCDE onto the activated sensor. The enzyme loaded AR2G sensors are then immersed in titre wells containing shell proteins. The association and dissociation kinetics are recorded. The sensors are regenerated using 2M NaCl. Global fitting of the association and dissociation kinetics is performed using Data Analysis HT 9.0.0.33 software provided with the OctateK2 instrument.

#### Aggregation kinetics experiment

0.5 mg/ml of PduCDE is mixed with 1mg/ml of shell proteins (only buffer in control sample) and incubated at 4°C for 1 hr. For aggregation studies, protein samples are maintained at 45°C in Spectrofluorimeter FS5 (Edinburgh Instruments, UK). The samples ate excited at 360 nm, band width of 1 nm and scattered light at 360 nm (band width of 1 nm) are recorded as a function of time.

#### Fluorescence anisotropy

1 mg/ml of PduBB’ is labeled with Acrylodan in 5:1 (protein: dye in μM) ratio. The samples are incubated overnight at 4°C. The dye labelled protein is separated from free dye using PD10 columns. The presence of dye is confirmed using UV-Vis spectrophotometer and the presence of protein is confirmed using and Bradford assay. The shell protein and enzyme are mixed and incubated at 4°C for 1 hour. Time resolved fluorescence anisotropy measurements are carried out using time correlated single photon counting (TCSPC) (Horiba, Japan) instrument equipped with 375 nm laser (Nd:YAG). 300ul of protein samples are used in the experiment. Fluorescence emission is collected at 485 nm emission wavelength and at 0° (parallel) 90° (perpendicular) angle with respect to excitation light. Perpendicular component of the emission is corrected using the G-factor. G-factor was determined independently for the free dye. Instrument response function was found to be ∼ 180 ps. Anisotropy decays are fitted using bi-exponential decay model:

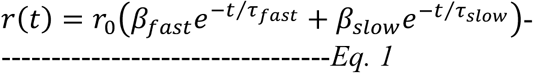

Here r_0_ is the intrinsic fluorescence anisotropy, τ_fast_ and τ_slow_ are faster and slower rotational correlation time and β_fast_ and β_slow_ are amplitudes of faster and slower rotational correlation time respectively.

#### Molecular docking

The modelled PduB’ and PduB are docked with Diol dehydratase (1DIO) using ClusPro server (https://cluspro.bu.edu/login.php). We select models with balanced electrostatic, hydrophobic and van der walls interactions. The docked model with the maximum number of clusters is selected for further analysis. The models are visualized in Pymol software (41). PDB sum server (http://www.ebi.ac.uk/pdbsum) (42) is used for identifying residual interactions. Peptide docking is performed using PIPER-Flex PepDock (http://piperfpd.furmanlab.cs.huji.ac.il/). In case of docking between N-terminal of PduD with PduB trimer, 9 out of 10 models show binding of the terminal residues of PduD to the concave side of PduB trimer. The docked model with the maximum interface score is selected for analysis. The model showing interaction between the terminal residues of PduD and convex side of PduB trimer (only 1 out of 10 models) is also selected for analysis. PDB sum server is used for identifying residual interactions.

#### Divergence Timeline

The timeline of divergence of thermophiles and Salmonella is generated using Time Tree webserver (43).

## Acknowledgments

S.S acknowledges financial support by SERB-DST, Govt. of India (Grant EMR/2015/000746 and SR/NM/NB-1082/2017). G.K and NKB thank INST, Mohali for fellowship. The authors thank Dr. Sabyasachi Rakshit, (IISER, Mohali) for helpful suggestions and support.

## Competing interest

The authors declare that there are no competing interests associated with the manuscript.

## Author Contributions

S.S. conceived the idea. G.K. and S.S. planned and designed the experiments. G.K., N.K.B. and J.P.H. executed the experiments. G.K. and S.S. wrote the paper. All the authors discussed the results and contributed to the present manuscript.

## The abbreviations used are

BMC: Bacterial microcompartments;
MCP: Microcompartments;
BLAST: Basic Local Alignment Search Tool;
ANS: 8-Anilino-1-npthalenesulfonic acid.

